# Approaches for calculating solvation free energies and enthalpies demonstrated with an update of the FreeSolv database

**DOI:** 10.1101/104281

**Authors:** Guilherme Duarte Ramos Matos, Daisy Y. Kyu, Hannes H. Loeffler, John D. Chodera, Michael R. Shirts, David L. Mobley

**Author notes:** University of California, Irvine (Guilherme Duarte Ramos Matos). University of California, Irvine (Daisy Y. Kyu). STFC Daresbury, Warrington, WA4 4AD, United Kingdom. University of Colorado, Boulder (Michael R. Shirts). University of California, Irvine (David L. Mobley).

## Abstract

Solvation free energies can now be calculated precisely from molecular simulations, providing a valuable test of the energy functions underlying these simulations. Here, we briefly review “alchemical” approaches for calculating the solvation free energies of small, neutral organic molecules from molecular simulations, and illustrate by applying them to calculate aqueous solvation free energies (hydration free energies). These approaches use a non-physical pathway to compute free energy differences from a simulation or set of simulations and appear to be a particularly robust and general-purpose approach for this task. We also present an update (version 0.5) to our FreeSolv database of experimental and calculated hydration free energies of neutral compounds and provide input files in formats for several simulation packages. This revision to FreeSolv provides calculated values generated with a single protocol and software version, rather than the heterogeneous protocols used in the prior version of the database. We also further update the database to provide calculated enthalpies and entropies of hydration and some experimental enthalpies and entropies, as well as electrostatic and nonpolar components of solvation free energies.

## Introduction

Solvation free energies give the free energy change associated with the transfer of a molecule between ideal gas and solvent at a certain temperature and pressure. While solvation free energies (Δ*G^solv^*) in general, and hydration free energies (Δ*G^hyd^*, solvation in water) in particular might not seem to have far reaching implications, in fact, researchers in diverse areas can benefit from their prediction, because such solvation free energies are related to a broad range of physical properties such as infinite dilution activity coefficients, Henry’s law constants, solubilities, and distribution of chemical species between immiscible solvents or different phases. Solvation free energies (*DeltaG^solv^*)—especially hydration free energies (Δ*G^hyd^*, aqueous solvation free energies)—are related to a broad range of physical properties such as infinite dilution activity coefficients, Henry?s law constants, solubilities, and distribution of chemical species between immiscible solvents or different phases.

Solvation free energies are differences in thermodynamic potentials which describe the relative populations of a chemical species in solution and gas phase at equilibrium.^1,2^ In the thermodynamic limit in the solvated phase and the ideal gas limit in the gas phase, Δ*G^solv^* of component *i* is equal to *μ_i,solv_ – μ_i,gas_*, the difference in chemical potentials in the two phases. In the additional limit of one molecule of component *i* at infinite dilution, these become the infinite dilution excess chemical potentials in the respective solvents.

Solvation free energies not only tell us how much a molecule prefers one phase over another, but they also can provide insight into how solvent behaves in different environments. For example, water solvates molecules of opposite polarity differently, due to its inherent asymmetry,^3^ surfaces also have asymmetric effects on ion pairing which depend on the curvature of the surface,^4^ and molecular geometry and chemical environment affects hydrophobic solvation.^5^ Although they can be difficult to measure experimentally, Δ*G^solv^* and Δ*G^hyd^* can be calculated to a precision better than 0.4 kJ·mol^−1^, even with a relatively modest investment of simulation time, for relatively diverse small neutral molecules^6^ such as those seen in the FreeSolv database of hydration free energies^7^ and in recent blind challenges such as the Statistical Assessment of the Modeling of Protein and Ligands (SAMPL) challenges. These challenges aim to improve the quality of predictive computational tools in drug design,^1,6,8–21^ and have leveraged solvation free energies to help drive improvements in modeling.

Since the solvation free energy of neutral compounds is an aggregate measure of many competing interactions and entropic effects that can span many kJ/mol, comparison of computed solvation free energies to experiment has proven to be an exacting test of force field quality that has been useful in revealing deficiencies in small molecule force fields.^3,22,23^ The relative ease by which solvation free energies can be calculated – as opposed to protein-ligand binding free energies, which are fraught with a variety of sampling issues – also makes them attractive for this purpose^a^. For instance, SAMPL has frequently (in SAMPL1 through SAMPL4) included blind predictions of hydration free energies in particular.^1,8–14^ However, to our knowledge, no laboratories are currently measuring hydration free energies, leading the field to search for other simple physical properties that can be rapidly computed – such as relative solubilities,^24^ distribution coefficients,^25^ and solvation free energies in organic solvents^26^ – as a tool to assess and improve small molecule force fields. In computational chemistry, hydration free energies are of particular importance because they are frequently used in force field parameterization^26–29^ and in the testing of free energy methods and force fields.^1,8–14,30–37^

Solvation free energies are often calculated by alchemical free energy methods,^38^ which simulate a series of alchemical intermediates to compute the free energy of transferring a solute from solution to gas phase (as here) or vise versa. This alchemical path provides an efficient way to move the solute from solution to the gas phase by perturbing its interactions in a non-physical way. Since free energy is path-independent, this non-physical process still yields the free energy change for transfer of the solute from solvent to gas.^38,39^ The path is formed by constructing intermediate states with interactions that modulate between the end states of interest, with the variable *λ* parameterizing progress along the path. A particularly efficient set of intermediate states uses a two step process, first turning off the van der Waals interactions using one parameter *λ_v_*, and another turning off the electrostatic interactions using a second *λ_e_*. Here, we compute the free energy change to transition between each pair of *λ* values, and the overall free energy change is the sum of these pairwise differences.

While other approaches have been used to calculate solvation free energies,^40^ alchemical free energy calculations using explicit solvent have become a mainstream approach,^41,42^ in part because of their formal rigor. Alternative approaches include implicit solvent models,^34–37,43^ which yield Δ*G^hyd^* but do not take into consideration solvent configuration around the solute, and Monte Carlo based approaches using the Gibbs ensemble^44–50^ and expanded ensemble,^51^ though these are most commonly used for molecules that are particularly small and/or rigid.

## Hydration and solvation free energies have a range of applications

The activity coefficient γ_*i*_ of a solute species *i* can be calculated from Δ*G^solv^*:

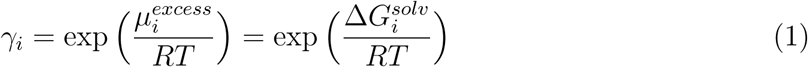

Where *μ*_*i*_^excess^ is the excess chemical potential of *i* and is equal to Δ*G^solv^_i_* in the ideal gas limit of the vapor phase, *R* is the universal gas constant and *T*, the absolute temperature. For instance, solvation free energies are used to estimate infinite dilution activity coefficients 
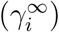
 in many solvents by using a single molecule of solute *i*.^52–58^ Experimental results obtained from gas chromatography^59,60^ can be compared to 
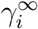
 obtained from Δ*G^solv^* to further test models and methodologies that use these free energy calculations.

Solubility prediction is another field where Δ*G^solv/hyd^* prediction can have great value. One methodology computes the free energy of solubility by computing both the sublimation free energy (from solid to gas) and hydration free energy (from gas to water).^61^ Another way to predict molecular solid solubilities depends on excess chemical potential calculations. The chemical potential, *μ*, of a species is calculated at different concentrations to build the concentration-dependent chemical potential curve of solutions^62–65^ in order to discover phase equilibrium conditions. Free energies of solvation in pure melts and pure amorphous matter have been used to find upper bounds for solubilities given that most drug-like compounds have crystal polymorphs.^66–69^ Relative solubilities of a given chemical species between different solvents can also be assessed with these calculations.^24,70^ Henry’s law solubility constants^71^ and solubilities in supercritical fluids^72^ can also be predicted using solvation free energies.

The latest SAMPL challenge, SAMPL5, included blind prediction of distribution coefficients between cyclohexane and water for 53 solutes.^32,33,73,74^ Distribution and partition coefficients are important properties for toxicology and pharmacology because they play a major part in predicting absorption and distribution of a substance in different tissues.^75^ Partition coefficients – which which are the distribution coefficients of the neutral form of a compound – can be estimated from the difference between solvation free energies of the chemical species in two different solvents,^21^ as shown in equation 2:

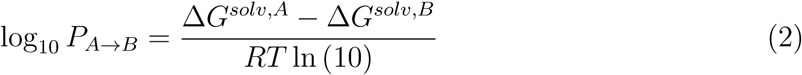

Where Δ*G^solv,A^* and Δ*G^solv,B^* are the solvation free energies of a molecule in solvents *A* and *B*, respectively. While in principle, the calculation could be done by transferring the solute between phases, in many software implementations it is more straightforward to simply compute the solvation free energy in each phase separately, or the free energy of removing the solute from each phase. Thus, solvation free energy calculations have found relatively widespread application in calculating partition coefficients, including in SAMPL5.^15–21^ Hydration free energies themselves are valuable quantities in drug design^42,76^ and can be used to understand the impact desolvation of the ligand has in the binding process^77,78^ or can be utilized as QSAR descriptors.^79^

## Theory and practical aspects of alchemical calculations

Solvation free energies can be calculated in various ways. In this paper we focus in alchemical free energy calculations, which have been one of the most consistently reliable methods in recent applications such as the SAMPL series of challenges.^1,8–14,25^ Consider a pair of end states *A* and *B*, and their respective Hamiltonians *H*_*A*_(**q**, **p**, ***λ***) and H_B_(q, p, ***λ***).

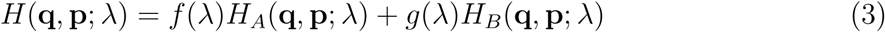

Where *f*(*λ*) and *g*(*λ*) are functions of lambda used to mix the Hamiltonians, typically set such that *H* = *H*_*A*_ at *λ*= 0. With *H*(**q**, **p**, *λ*) we can calculate the free energy difference between *A* and *B*:

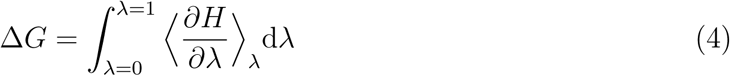

This method, called thermodynamic integration (TI),^80^ performs similarly to more robust methods when the integrand is smooth.^81–83^ However, it can break down when the integrand is not smooth, and it can be difficult to capture numerical integration errors in resulting uncertainty estimates.

Exponential averaging (EXP), also known as Free Energy Perturbation (FEP), was introduced by Zwanzig.^84^ In this method, the free energy difference between two states *A* and *B* is given by:

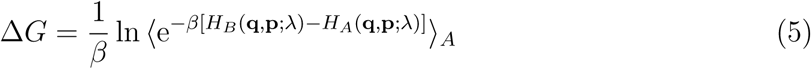

Where *β* = (*k*_*B*_*T*)^−1^. Although equation 5 is exact in the limit of large numbers of samples, EXP is inefficient and prone to shot noise and other large biases when configurations sampled in one state are very unlikely to be found in the other state, and vice-versa. The probability that describes this likelihood is called the phase-space overlap between the two states. EXP convergence is far from ideal, requiring states to have sizable phase-space overlap with one another.^38,82,85^ Thus, addition of intermediate states (with values of *λ* between 0 and 1) can improve overlap dramatically and thus the quality of the final result.^86^ Another issue is an asymmetric bias depending on which direction the free energy difference extrapolation is performed,^87,88^ so other analysis methods are now preferred.^38^ In the limit of adequate sampling, EXP converges to the same free energy value in both directions, but there are other ways to calculate free energies more efficiently.

An alternate method, Bennett’s acceptance ratio (BAR), uses the information from both directions to derive the following relationship:

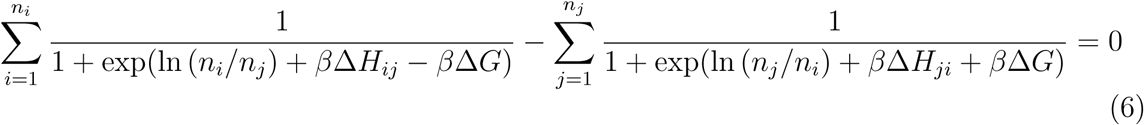

Where *n_i_* and *n_j_* are the number of samples between each state and Δ*H_ij_* is the enthalpy difference between these states.

This expression minimizes the free energy variance^89^ and makes BAR much more efficient than EXP.^87,88^ The Multistate Bennett acceptance ratio (MBAR) is an extension of BAR that considers the overlap between a given state and all the others in the path between the end states.^90^ BAR and MBAR perform similarly when the spacing between intermediate states is moderate^83^ and therefore only neighboring states have phase-space overlap. WHAM^91,92^ is essentially an approximation to MBAR, and thus also gives very similar results when carefully done with appropriately small bins. MBAR performs consistently well, and indeed is perhaps the most consistently well-performing free energy estimator, ^83^ thus we recommend it as the analysis method of choice whenever possible. In many cases, TI can perform as well as BAR or MBAR if the integrand is smooth.^81,82^ EXP should generally be avoided due to its asymmetric bias and sensitivity to the tails of the distribution.^87^

### Choice of alchemical pathway

Alchemical free energy calculations were given this name because the pathway involves unphysical changes to the atomic identities, such as to the interactions between components.^42,93^ Solvation free energy calculations can use several different approaches to modulating interactions. One approach, called *decoupling*, modulates only the interactions between the solute and its surroundings, retaining internal interactions (which is the approach we use here). An alternative approach, called *annihilation*, removes internal non-bonded interactions within the solute as well as those with the surroundings. Mixtures of the two approaches are also possible, such as annihilating internal electrostatic interactions while decoupling non-polar interactions. Here, three main states are considered: a single, non-interacting molecule of the solute in a box of solvent; the solute molecule that interacts with its surroundings through nonpolar (dispersion and exclusion) forces; and a fully interacting system, in which solvent molecules interact with the solute molecule through both electrostatic and nonpolar (dispersion and exclusion) forces. Simulations are then conducted over a series of intermediates connecting these states: going through a phase which changes electrostatic interactions only, and another phase which modifies van der Waals interactions only (figure 1). Each of these intermediates has high configuration space overlap with at least neighboring states, allowing precise calculation of free energy differences.^94–97^

**Figure 1:**
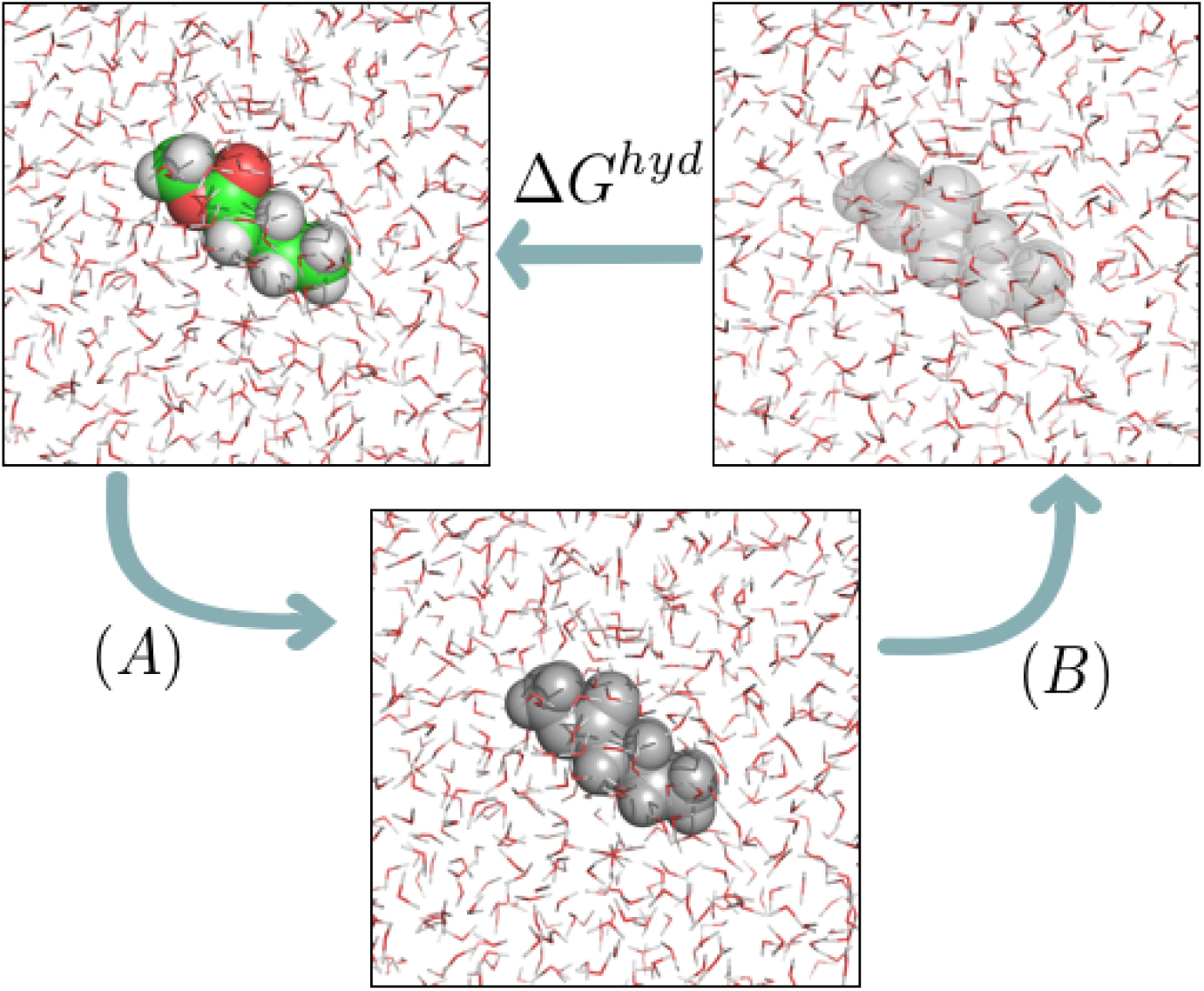
Thermodynamic cycle used to calculate hydration free energies (or, more generally, solvation free energies). In (*A*), we have states in which charge-charge interactions between the solute and its environment are progressively turned off. In (*B*) dispersion interactions between solute and water are progressively turned off. Colored atoms (green for carbon, red for oxygen, white for hydrogen) have electrostatic and nonpolar interactions with the environment; gray atoms retain only nonpolar interactions; and transparent atoms have no interactions with their environment (and thus represent the solute in vacuum).

The most straightforward way to switch between states is the linear pathway

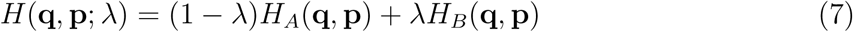

but this pathway is in general problematic for solvation of all but the smallest molecules. This is because repulsive and forces are often handled by a 1/*r*^12^ term (such as in the Lennard-Jones functional form) which leads to non-integrable singularities in 〈ðH/ðλ〉 at terminal *λ* values due to sudden changes in the potential at small *r*. This is a not a problem which is specific to TI; rather, this issue can still result in numerical instabilities or large errors in calculated free energies even with other analysis approaches.^42,98,99^ Thus, more complicated *λ* pathways are required, such as soft-core potentials, which should in general be used to avoid such numerical problems.^94,98,99^ A common soft-core form for Lennard-Jones potential between two particles *i* and *j* is:

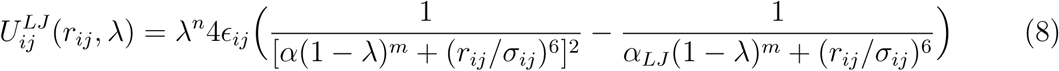

Where *ϵ*_*ij*_ and *σ*_*ij*_ are the Lennard-Jones parameters and *α* is a positive constant which should typically be set to 0.5.^99,100^ The exponents *m* and *n* are most efficient at *n* = 1 and *m* = 1 but other values have been used too.^96,99–101^ Improvements have been achieved by new soft-core functions that ease the problem with additional minima within the formulation of the original soft-core potential,^102^ and alternate potentials that construct near optimal paths for alchemical simulations.^103^ Linear basis functions can be used as an alternative to soft-core potentials that approaches the minimum variance possible over all pair potentials;^97,104^ these can also enhance the efficiency of alchemical calculations.”

The use of soft-core potentials promotes better convergence in many circumstances, and provides much lower variance free energy estimates given a fixed amount of simulation time,^94,96,98,100,103^ thus their use is highly recommended for successful free energy calculations. Without soft-core potentials, convergence is much more difficult or nearly impossible to achieve in many types of solvation free energy calculations.

### Considerations for successful alchemical calculations

The accuracy of these calculations is affected by at least three factors:^105,106^ Is our sampling representative and adequate? Is the free energy estimator good enough? Is the force field adequate for the system? For solvation free energies of small molecules in solvents with relatively fast dynamics, such as water, sampling is typically adequate with a few nanoseconds of dynamics per lambda window (at least for relatively rigid solutes), and the free energy estimators above are robust when applied carefully.

However, when designing new studies, it is still important to choose robustly performing estimators and ensure adequate sampling. As discussed above, we recommend MBAR as the best and most reliable general-purpose estimator.^83^ Sampling remains a critical issue,^105,107^ both as the solute size and flexibility grows and as solvent dynamics or environment become heterogeneous, for example, for solvation free energies in octanol which can form local clusters of hydrophilic and hydrophobic sites,^21^ or in mixed solvents.^25^

## We updated FreeSolv, the free community solvation free energy database

FreeSolv^7^ is a hydration free energy database for neutral^b^ compounds that contains experimental and calculated hydration free energy values, SMILES strings, PubChem compound IDs, IUPAC names, and now (as of version 0.5, presented in this work) calculated enthalpies and entropies of hydration of 643 small organic molecules. Since experimental and calculated hydration free energies, Δ*G^hyd^*, can be computed quite precisely for quantitative comparison, FreeSolv can provide information for force field development,^26–29^ and can assist the testing of new solvation free energy methods.^108,109^ While calculated hydration free energies for all compounds have been in FreeSolv since the database was constructed,^7^ previous values had been calculated with somewhat heterogeneous protocols in a variety of different studies spread over roughly 10 years.^2,6,11,13,23,41,110,111^ In this work, we have updated FreeSolv by repeating all of the calculations under a single unified protocol, and additionally obtained enthalpies and entropies of hydration.

We obtained FreeSolv’s calculated hydration free energies using alchemical free energy calculations, connecting the end states (corresponding to the solute in vacuum and in solution) via a *λ* path with 20 intermediate states (full details in SI). The first five states corresponded to changes in electrostatic interactions, while the last 15 modified the Lennard-Jones terms in the potential. This separation allows electrostatic interactions to be changed linearly, and soft-core potentials to be used only when changing non-polar interactions.^97^ We ran 5 nanoseconds of Langevin dynamics per state with 2 femtosecond time steps in GRO-MACS 4.6.7^112–117^ at 298.15K. Pressure was maintained at 1 atm by the Parrinello-Rahman barostat.^118^ Full details can be found in the supporting materials.

Input files for version 0.5 of FreeSolv were constructed from scratch from the isomeric SMILES strings for the compounds which are deposited in the database. From these SMILES strings, we used the OpenEye Python toolkits^119–121^ to generate molecular structures and assign AM1-BCC partial charges,^122,123^ then charged mol2 files were written out. The AMBER Antechamber package was then used to to assign parameters from the GAFF^20^ small molecule force field (version 1.7), and these were then converted to GROMACS format and solvated with the TIP3P water model.^124^ The script which performs the setup and regenerates all input and molecular structure files in the database is available in the scripts directory of FreeSolv and provides full details. Following the calculations, MBAR hydration free energies were obtained using alchemical-analysis.py.^93^ Additional details can be found in the supporting material.

Computed hydration free energies are compared with experiment in figure 2.

In the calculations described in this study, we found an average error of 1.3±0.3 kJ·mol^−1^, RMS error of 6.4 ± 0.3 kJ·mol^−1^, average absolute error of 4.7 ± 0.2 kJ·mol^−1^, Kendall *τ* of 0.80±0.01, and Pearson *R* of 0.933±0.008, comparable to those in the original FreeSolv set,^7^ though some individual compounds have reasonably significant discrepancies (see SI). This level of accuracy is consistent with what is often seen from classical fixed-charge force fields, which typically yield RMS errors around 4-8 kJ/mol in computed hydration free energies.^42^

**Figure 2:**
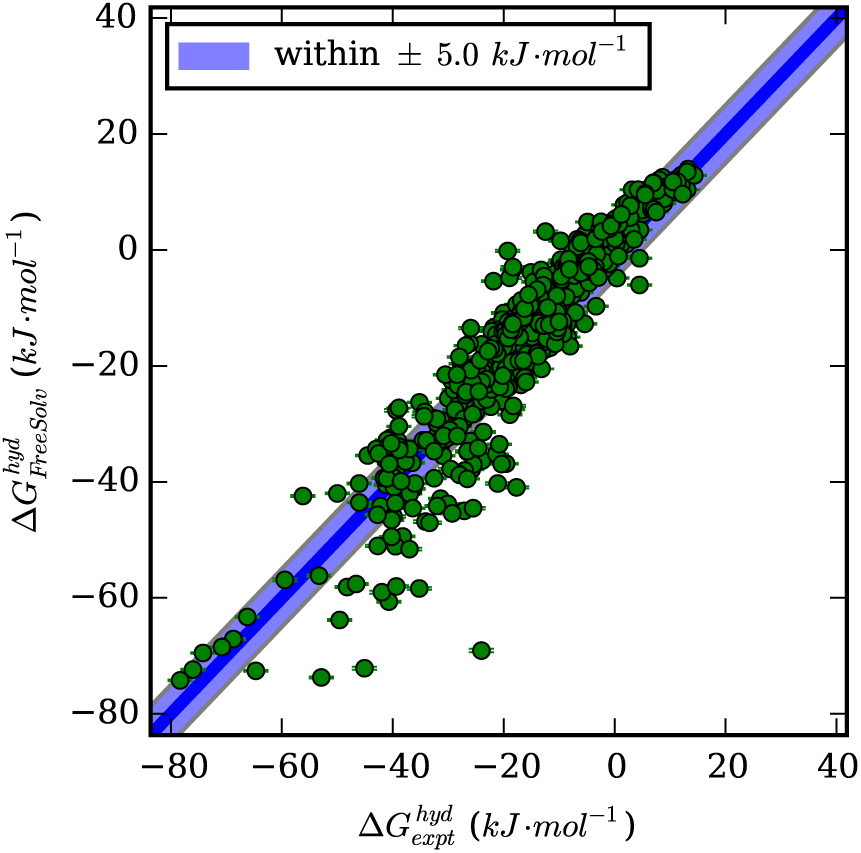
Calculated versus experimental hydration free energies for the compounds in Free-Solv version 0.5.

In addition to experimental and calculated values, FreeSolv now includes the free energy of decoupling the solute-solvent electrostatic interactions (Δ*G^q^*) and the free energy of decoupling the nonpolar interactions in water (Δ*G^vdW^*) (available at github.com/mobleylab/ FreeSolv). These quantities have been used for various purposes, including to assist in the study, development, and test of implicit solvent models.^125,126^ However, it is important to remember that these components come from our particular decomposition of the free energy,^127–130^ and are not state functions; other decompositions are possible, so considerable care needs to be taken in interpreting these components. For example, annihilation rather than decoupling of Coulomb interactions would result in somewhat different decompositions due to electrostatics-induced conformational differences while van der Waals interactions are being decoupled.

In addition to hydration free energies, we have also computed enthalpies (Δ*H^hyd^*) and entropies of hydration (Δ*S^hyd^*), and have added these to the database. Enthalpies of transfer, due to their larger dynamic range and lack of compensating entropic effects, are generally more sensitive to force field parameters than free energies,^131–133^ and thus can be sensitive probes of force field accuracy, providing an additional point of comparison to experiments. While only a few hydration enthalpies are available experimentally, there are a sufficient number to note that significant discrepancies between experiment and computed values exist for some compounds (SI Figure 2 and SI Table 1). There seems to be little correlation between compounds which have accurate hydration free energies versus which have accurate hydration enthalpies; for example, the calculated hydration free energy of benzene is within error of the experimental value, but the enthalpy is off by approximately 12 kJ/mol. In contrast, the hydration free energy of cyclohexanol is off by more than 5 kJ/mol but the enthalpy is within error of the experimental value. Thus, clearly these quantities yield different information. We postulate that, should more experimental enthalpies of hydration become available, this information may provide additional insight into force field deficiencies.

To compute hydration free energies via simulation, we used a difference in potential energies between a water box solvating the compound and a neat water box with the compound removed to vacuum:

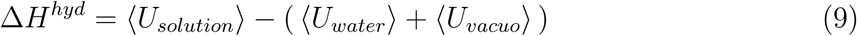

Here, ⟨*U_solution_*⟩ is the internal energy of the solution (containing the solute); ⟨*U_water_*⟩ is the internal energy of a box of the same number of water molecules (under the same conditions) without the solute; and ⟨*U_vacuo_*⟩ is the internal energy of the solute molecule alone in vacuum. We have neglected the pressure-volume contribution to the enthalpies, *P*Δ*V*, since for solutes of this size, the contribution is much smaller than our typical uncertainties of ≈ 2.9 kJ · mol ^−1^;^131^ at larger pressures or for larger solutes than in this set, this term could become significant. Notably, this scheme also omits other contributions that may be relevant in making direct comparison with experimental enthalpies of hydration, including contributions from the cost of polarizing the molecule from vacuum to solvated phase charges (relevant to fixed-charge force fields), corrections to the vibrational modes due to the quantum chemical nature of real solutes, nonideality of the gas phase, and the fact that the simulation of the liquid is carried out at atmospheric pressure rather than at the vapor pressure of the gas phase; for a review of these contributions, see.^134^ We note that other groups have also omitted these contributions, which still await a thorough assessment of relative magnitude for small molecule hydration enthalpies.^131^

To calculate the average potential energies, we ran 60 nanosecond Langevin dynamics simulations, with two femtosecond timesteps at 298.15 K and 1 atm in water and *in vacuo* for each molecule in the database. These long simulations were necessary to reduce error bars on the computed enthalpies to levels around 2.9 kJ · mol ^−1^, roughly the level of typical thermal energy ( 1 *k_B_T*) as done in, for example, host-guest binding calculations.^132^ In order to obtain consistent results, we used simulation boxes with 1; 309 water molecules and one solute molecule. The same system parameters and water model were used as in the free energy calculations. Hydration entropies are calculated via the equation:

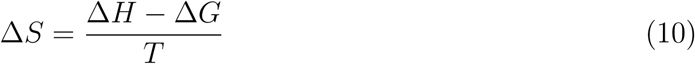

With Δ*G* and Δ*H* calculated as described previously.

Calculated hydration enthalpies exhibit some correlation with calculated hydration free energies, but the correlation is not perfect, indicating that enthalpies can indeed provide additional constraints on the force field.^132^ The Kendall and *τ* the Pearson *R* between the calculated Δ*H^hyd^* and Δ*G^hyd^* respectively were 0.76 ± 0.02 and 0.943 ± 0.005 (see supplementary information).

Our conclusion that enthalpies can provide an additional constraint on the force field is further supported by comparison to experimental data. Specifically, 11 experimental hydration enthalpies and entropies from ORCHYD, a database of experimental hydration properties,^135^ were added to FreeSolv. Calculated and experimental enthalpies have a Kendall *τ* of 0.77 ± 0.05, and a Pearson *R* of 0.87 ± 0.03 (see SI). These values indicate that the computed hydration free energies are relatively predictive of experimental values, though there is also clear room for improvement. Calculated hydration enthalpies and their experimental counterparts show significant differences that are not observed in the plot of experimental versus calculated free energies of the same 11 compounds, suggesting (as in previous studies^131^) that enthalpies provide additional information on the thermodynamics and constraints on the force field (though as noted above, additional enthalpy corrections may be needed^134^). More details can be found in the supporting information.

We also partitioned the hydration enthalpy, Δ*H*, into two components: a solvent interaction term and a conformational change term, 
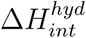
 and 
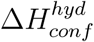
 respectively, in order to understand how much the solvation enthalpy is influenced by the solute conformation, and how much solute conformation is modulated by solvation. We obtained the solvent interaction component by taking the average energy of the solute in water and subtracting off the solute internal energy and the energy of a corresponding box of pure water, leaving only the enthalpy change due to changing solute-solvent interactions and solvent reorganization:

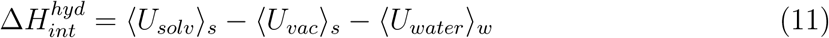

where 〈*U _solv_*〉*s* is the average potential energy over the original solvated trajectory, 〈*U_vac_*〉*s* is the average potential energy of the solute molecule in the solvated trajectory after removing its water molecules, and 〈*U_water_*〉*_w_* is the average potential of a simulation with an equivalent box with pure water. 
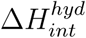
 thus corresponds to the change in solvation enthalpy due to transferring a solute molecule from vacuum to water with a *fixed set of configurations* (as given by the solvated trajectory) – i.e., it treats the solute as if there is no conformational change going from gas to water, so it includes only changes in solvent structure and solute-solvent interactions.

The conformational change component of the enthalpy is obtained by taking the change in solute internal energy on going from gas to water, which we can evaluate as follows:

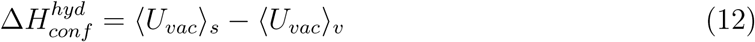

where 〈*U_vac_*〉*_v_* is the potential energy of the solute molecule in vacuum evaluated from the trajectory run in vacuum, and 〈*U*_*vac*_〉*_s_* is the potential energy of the solute molecule in vacuum evaluated from the trajectory run in solvent (after stripping the solvent molecules). 
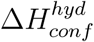
 thus gives to the enthalpy change due to solute conformational changes on solvation; these occur because interactions with water can stabilize configurations that are not common in vacuum. If a compound’s distribution of configurations is unchanged on transfer to solvent, 
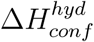
 will be zero. It can trivially be verified that these components still sum to the total enthalpy change:

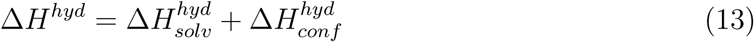

These components, while certainly not a unique decomposition of the total enthalpy, do provide a way to intuitively understand one important set of contributions to the enthalpy of hydration in a way which provides some insight into changes undergone by the solute and environment. For example, solutes which undergo significant conformational changes on solvation may tend to have a large change in the conformational component of the hydration enthalpy (fig. 3). This happens because solutes that make hydrogen bonds with water or have strong internal electrostatic interactions in the gas phase can assume conformations that were energetically unfavorable *in vacuo* when solvated.

Our intention is that FreeSolv serve as an updateable, extensible community resource. While it already covers a large number of molecules, we would be delighted to include input files and calculated values from other force fields and/or methods so it can further serve as a benchmark of methods, simulation packages, and so on. Additionally, while hydration free energy data is not abundant, certainly at least some data is available that is not presently included in FreeSolv, so community contributions of experimental data with references will be appreciated. Additional curation of the experimental data already present is likely needed – for example, much of the experimental data still needs to be tracked back to its original source material rather than literature compilations of data which are currently cited. FreeSolv is available on GitHub at http://github.com/mobleylab/FreeSolv and contributions are welcomed there.

**Figure 3:**
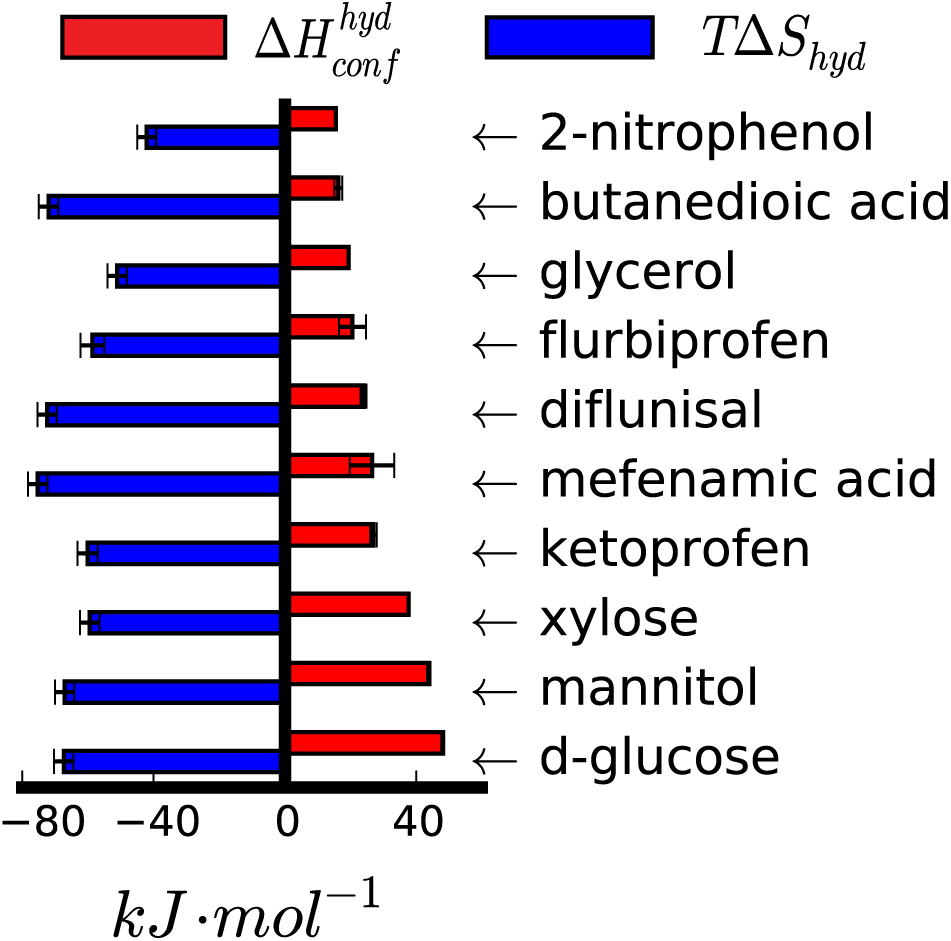
Conformational enthalpies and associated entropies of compounds with highest 
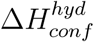
.

## Conclusions

Solvation free energies have been the subject of considerable scientific interest for many years because they are related to a large number of physical properties. Here, we have provided a short review of alchemical methods for computing solvation free energies of small organic molecules, and discussed their application to hydration free energies. Solvation free energies for such molecules can be calculated precisely and effectively using alchemical free energy calculations, as described here, preferably using MBAR as the free energy estimator.

We also introduced an update to FreeSolv^7^ (v0.5), a database of calculated and experimental hydration free energies, enthalpies and entropies. The database was designed to be easily incorporated into automated workflows: we provide IUPAC names, PubChem compound IDs and SMILES strings, as well as topology and coordinate files, but additional data is welcome. Additionally, we provide calculated and experimental free energy values that can be used to assist method and force field development. Unfortunately, experimental hydration enthalpies and entropies are not available for every compound.

Calculated free energies show reasonable agreement with experimental values (fig. 2) with an RMS error around 6 kJ· mol^−1^ and an average error close to 1 kJ· mol^−1^. With the aid of ORCHYD,^135^ we were able to extend FreeSolv to contain experimental hydration enthalpies for a few (11) compounds for the first time. We observe significant errors for hydration enthalpies that are much larger than those for hydration free energies, so further investigation will be needed. This result also suggests that enthalpies can be used as additional constraints in force field development.

We believe that this update of FreeSolv will assist future efforts in force field development and development and testing of new methods. We also hope that FreeSolv’s new features help serve the scientific community, and provide a valuable resource the community will help extend.

## Acknowledgement

DLM and GDRM appreciate the financial support from the National Science Foundation (CHE 1352608), and computing support from the UCI GreenPlanet cluster, supported in part by NSF Grant CHE-0840513. GDRM appreciates support from the Brazilian agency CAPES - Science without Borders program (BEX 3932-13-3). JDC acknowledges partial support from NIH grant P30 CA008748. HHL is supported through an EPSRC provided SLA, funding the core support of CCPBioSim. CCPBioSim is the Collaborative Computational Project for Biomolecular Simulation funded by EPSRC grants EP/J010588/1 and EP/M022609/1. We particularly appreciate Kyle Beauchamp (Counsyl, South San Francisco, CA) and Lee-Ping Wang (UC Davis) for input on curation of the FreeSolv database. We also thank Gaetano Calabrò and Caitlin Bannan for their support and assistance, and sharing of knowledge.

DLM is a member of the Scientific Advisory Board for OpenEye Scientific Software. JDC is a member of the Scientific Advisory Board for Schrödinger, LLC.

## Supporting Information Available

The following files are available free of charge.

- Supplement.pdf: document containing correlation plots between calculated free energies, enthalpies and entropies.
- GROMACS 4.6.7 .mdp files: GROMACS input files containing all the details of the simulations.
- FreeSolv can be obtained free of charge at http://github.com/mobleylab/FreeSolv. This material is available free of charge via the Internet at http://pubs.acs.org/.

## Footnotes

a But see the Supporting Information for how protonation state/tautomer challenges may apply here, as in protein-ligand binding.

b For additional discussion of why the focus is on neutral compounds, see the Supporting Information.

---

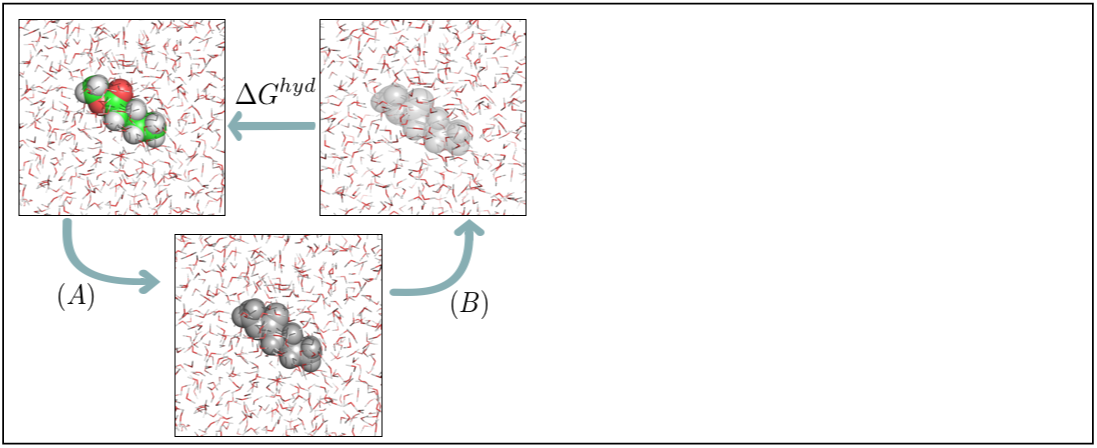
Graphical TOC Entry

